# Molecular mechanism of NAD^+^ binding to the Nudix homology domains of DBC1

**DOI:** 10.1101/2023.10.27.564493

**Authors:** Liming Ou, Xuechen Zhao, Ivy (Jing) Wu, Zhiyuan Xiong, Zhi Ruan, Guangyu Zhou, Wen Chen

## Abstract

DBC1 (deleted in breast cancer 1) is a human nuclear protein that modulates the activities of various proteins. NAD^+^ (oxidized form of nicotinamide adenine dinucleotide) is thought to potentially bind to the Nudix homology domains (NHDs) of DBC1, thereby regulating DBC1-PARP1 [poly (adenosine diphosphate–ribose) polymerase] interactions, the modulation of which may restore DNA repair to protect against cancer, radiation, and aging. Therefore, our study comprehensively employed methods including NMR (Nuclear Magnetic Resonance), ITC (isothermal titration calorimetry), genetic mutation, and computer biology to thoroughly investigate the molecular mechanism of the binding interaction between NAD^+^ and its precursor NMN with the NHD domain of DBC1 (DBC1_354-396_). The results from NMR and ITC indicate that NAD^+^ likely interacts with DBC1_354-396_ through hydrogen bonding, with a binding affinity nearly twice that of NMN. The key binding sites are primarily E363 and D372. Molecular Docking further revealed the importance of conventional hydrogen bonds and carbon-hydrogen bonds in the binding process. These findings may lead to a better understanding of how NAD^+^ regulates the physiological functions of DBC1, thereby offering guiding principles for the development of targeted therapies and drug research focused on tumor diseases associated with DBC1.

## 1. Introduction

Deleted in breast cancer 1 (DBC1) is a nuclear protein, initially cloned from a human chromosome 8p21 region homozygously deleted in breast cancers[1]. It is one of the most abundant yet enigmatic proteins in mammals[2], with a conserved domain similar to Nudix hydrolases that hydrolyze nucleoside diphosphates but are catalytically inactive due to the absence of key catalytic residues[3, 4]. DBC1 was already known to bind and inhibit SIRT1, a NAD^+^-dependent deacetylase that is critically involved in stress responses, cellular metabolism, and aging[5, 6]. Of note however, DBC1 has recently emerged as an inhibitor of PARP1, another NAD^+^- dependent pleiotropic enzyme that plays a key role in DNA repair[7]. It is well known that DBC1 regulates a variety of targets by protein-protein interaction, contributing to various cellular processes including apoptosis, DNA repair, senescence, transcription, metabolism, circadian cycle, epigenetic regulation, cell proliferation, and tumorigenesis [6, 8-13]. However, to date information about how the interaction between DBC1 and its targets is regulated is scarce. Moreover, although the DBC1 gene localizes at a homozygously deleted region in breast cancer, its role in tumorigenesis remains unclear. It has been proposed to function either as a tumor suppressor or an oncogene[14, 15]. Therefore, further investigation is required to elucidate the role of DBC1 in cancers.

NAD^+^ (Nicotinamide adenine dinucleotide) is a crucial coenzyme involved in redox reactions, playing a central role in energy metabolism[16]. It also acts as a substrate for signaling by the PARPs [Poly (ADP-ribose) polymerases]] and the Sirtuins (SIRT1-7) in the regulation of DNA repair, cell survival, and circadian rhythms, among other functions[17-19]. Recent studies have shown that NAD^+^ has a third function of directly regulating protein-protein interactions, the modulation of which may protect against cancer, radiation, and aging[7]. Nowadays, the potential of NAD^+^ to anti-aging is of growing interest and is still being explored. Jun Li et al [7]revealed that NAD^+^ can bind directly to the NHD of DBC1, disintegrate the PARP1-DBC1 complex through competitive binding. This promotes DNA repair related to aging and radiation damage via PARP1 activity, and ultimately slowing aging. However, the precise mechanism of NAD^+^ binding to DBC1-NHD requires further investigation.

Endogenous and exogenous DNA damage accumulates over time, gradually impairing cellular function and accelerating the aging process and cancer[9, 20]. Emerging drugs that target DNA repair may delay age-related phenotypes or tumorigenesis. Deciphering the interaction sites between NAD^+^ and DBC1can provide a theoretical basis for understanding the molecular mechanism behind DNA repair, which may provide a reference for mining relevant targets and guiding the development of targeted drugs. Moreover, the research can provide valuable insight into the role of NAD^+^ in regulating DBC1 function, helping us better understand the role of DBC1 in tumorigenesis.

Accordingly, the aim of this work was to gain insight into the mechanisms of the interaction between NAD^+^ and DBC1-NHD fragment (residues 354-396, DBC1_354-396_) in vitro. We first expressed the DBC1_354-396_ protein with high purity and confirmed its activity using ^1^H-^15^N HSQC (Heteronuclear Single Quantum Coherence spectroscopy). Next, we assigned the protein backbone through triple resonance experiments (HNCA, CACB(CO)NH, combined with HNCACB). By performing NMR titration and quantifying the chemical shift changes of each amino acid residue, we identified which residues were involved in the interaction and the most significant interacting residues. And then we used isothermal titration calorimetry (ITC) to investigate the binding affinity of NAD^+^ and its precursor NMN to DBC1_354-396_. Subsequently, we mutated key amino acids in the protein and used ITC to determine if the mutated protein could still bind to NAD^+^. Finally, we employed molecular docking in computational biology to provide an in-depth elucidation of the intermolecular interactions and vividly explain the mechanism of binding.

## 2. Materials and methods

### 2.1 Material and chemicals

All chemicals were purchased from Sangon Biotech (China), with the following exceptions: Ni-NTA (Bio-Scale Mini Nuvia IMAC, 5ml) and gel filtration columns (Enrich SEC70) were purchased from Bio-Rad Laboratories, Inc (USA). NAD^+^ was procured from Solarbio (China), and NMN was sourced from Beyotime Biotechnology (China). ^15^NH4Cl and D-Glucose (U- 13C6, 99%) were obtained from Cambridge Isotope Laboratories, Inc (USA). DNA QuickMutation kit was purchased from Beyotime Biotechnology (China). The TEV protease was purified in-house by our research group.

### 2.2 Construction of DBC1_353-396_ vector

The DBC1-NHD fragment (residues 354-396) derived from the human DBC1 (Uniprot code: Q8N163) was synthesized and subsequently cloned into the pETM41 vector by GenScript Biotechnology Co., Ltd (Nanjing, China). The resulting plasmid encodes a fusion protein consisting of N-terminally hexahistidine-tagged MBP (maltose-binding protein) and DBC1, with a TEV protease recognition site in between. After purification and TEV protease cleavage, the final DBC1_354-396_ sequence will be <u>GAM</u>EEEAVLVGG**E**WSPSLDGL**D**PQADPQVLVRTAIRCAQAQTGIDL, with the three additional residues underlined representing remnants from the cloning process and the protease recognition site. Mutants of E363A/K and D372A/K were obtained using the DNA QuickMutation kit (Beyotime Biotechnology) and confirmed by DNA sequencing (mutations at positions E363 and D372 are indicated in bold).

### 2.3 Protein expression and purification

Each protein construct was expressed by cultivating transformed *E. coli* strain BL21 (DE3) cells in LB or M9 minimal media (when isotopic labeling was required). The cultures were grown at 37°C until OD600 reached 0.6, and then cooled to 18°C before induction with 0.4 mM isopropyl β-D-thiogalatopyranoside (IPTG). Protein was expressed at 18°C for 24 h. After expression, the cells were harvested through centrifugation at 5000×g and 4°C for 30 min.

The resulting pellet was then resuspended in lysis buffer (25 mM sodium phosphate, 100 mM NaCl, 20 mM imidazole, 2 mM PMSF, pH 7.4) plus a protease inhibitor cocktail (Biosharp). Cells were then lysed on ice by sonication using 250 W power with 2 cycles of 1.5s on and 1.5s off, each cycle lasts for 6 min. Lysate was separated into soluble fraction and inclusion bodies via centrifugation for 30 min at 37,500×g at 4°C. The soluble fraction was loaded onto a 5 mL Bio-Scale Mini Nuvia IMAC Ni-charged Cartridges (Bio-Rad) which was pre-equilibrated with 5 column volumes (CV) of equilibration buffer (25 mM sodium phosphate, 100 mM NaCl, 20 mM imidazole, pH 7.4). The column was then run on NGC 10 FPLC purification system (Bio-Rad) with a flow rate of 2 mL/min. The column was first washed with 6 CV of equilibration buffer, followed by washing with 6 CV of washing buffer (25 mM sodium phosphate, 100 mM NaCl, 50 mM imidazole, pH 7.4), and a subsequent linear gradient from 50 to 1000 mM imidazole in elution buffer (25 mM sodium phosphate, 100 mM NaCl, 1000 mM imidazole, pH 7.4) over 10 CV. The individual peaks were collected, and the fractions containing the MBP fusion protein were identified using SDS-PAGE. These fractions were dialyzed (3,000 MWCO) against exchange buffer for TEV protease digestion (50 mM Tris-HCl, 0.5 mM EDTA, 1 mM DTT, pH 8.0).

### 2.4 TEV protease digestion

We used a 1:100 enzyme to substrate ratio and conducted a 3-hours digestion at 30°C. The DBC1_354-396_ protein was collected using 5mL Bio-Scale Mini Nuvia IMAC Ni-charged Cartridges (Bio-Rad). Digestion efficacy was assessed with SDS-PAGE analysis. The flow-through containing DBC1_354-396_ protein was collected and concentrated using ultrafiltration tubes (MWCO=3000Da). The concentrated samples were loaded onto a gel filtration column (Enrich SEC70, 24 ml, Bio-Rad) for purification. The elution program consisted of a 30 mL volume at a flow rate of 0.75 mL/min, using a buffer composed of 25 mM HEPES, 150 mM NaCl, 2 mM Tcep, and pH 7.0 (NMR experiments: 25 mM MES, 2 mM Tcep, pH 6.5). Collected peak fractions were validated for protein purity by SDS-PAGE analysis and further concentrated using a 3000Da MWCO ultrafiltration tube.

### 2.5 Nuclear Magnetic Resonance (NMR) experiments

All NMR experiments were conducted at 25°C on Agilent 800 MHz spectrometer equipped with cryogenic probes. The NMR data were processed and analyzed using programs NMRpipe[21] and Sparky (T. D. Goddard and D. G. Kneller, SPARKY 3, University of California, San Francisc).

#### 2.5.1 HSQC of protein

For the acquisition of ^15^N-^1^H HSQC spectrum, 80 and 1000 points were taken in the N and H dimensions, respectively. Spectrum widths are 11 ppm in the H dimension and 22 ppm in the N dimension, while centering at 4.64 ppm in the H dimension and 119 ppm in the N dimension. 8 total scans were used.

#### 2.5.2 Backbone Assignment

A ^1^H-^13^C-^15^N triple-labeled protein sample was prepared for performing triple resonance experiments, including HNCA, CACB(CO)NH, HNCACB. By correlating the chemical shifts of Cαi-1, Cβi-1, Cαi, and Cβi atoms on the NMR spectra with the sequential order of protein amino acid residues, the identification and assignment of sequence-specific backbone atoms were achieved. The experimental parameters used for the assignment of the backbone in DBC1_354-396_ are shown in the table 1.

**Table 1.** NMR experiment parameters for DBC1_354-396_ backbone Assignment.

| Experimental Names | Spectral width<br>(H*C*N, ppm) | Center frequency<br>(H*C*N, ppm) | Complex points<br>(H*C*N) |
| --- | --- | --- | --- |
| <sup>1</sup>H-<sup>15</sup>N HSQC | 11 *22 | 4.64*119 | 2048*80 |
| <sup>1</sup>H-<sup>13</sup>C HSQC | 11*70 | 4.64*45 | 1000*120 |
| HNCACB | 11* 56*22 | 4.64*45*119 | 1000*70*41 |
| CACB(CO)NH | 11* 56*22 | 4.64*45*119 | 1000*56*41 |
| HNCA | 11* 30*22 | 4.64*54*119 | 1000*40*41 |

### 2.6 NMR titration

The protein sample concentration was 0.3 mM, and the NAD^+^ stock solution concentration was 75 mM, while the NMN stock solution concentration was 120 mM. According to the concentration ratio of protein to Ligands (1:0, 1:0.5, 1:1, 1:2, 1:4, 1:8, 1:16, 1:32), NAD^+^ and NMN were sequentially added to the protein, ensuring that the final volume of ligands did not exceed 1/20 to 1/10 of the total protein volume. ^15^N-^1^H HSQC spectra of protein samples under different ligand concentrations were recorded. By overlaying and comparing all ^15^N-^1^H HSQC spectra, and quantitatively analyzing the chemical shift perturbation (CSP) of each amino acid residue, it is easy to identify which residues are involved in interactions and which residue show the most significant effects[22]. The formula for quantitative chemical shift perturbation (CSP) analysis is as follows:

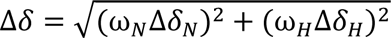

Δ*δ*_*H*_ represents the change in chemical shift of the 1H atom, and Δ*δ*_*N*_ represents the change in chemical shift of the 15N atom. The weighting factors for these changes are *ω*_*H*_ = 5.00 and *ω*_*N*_ = 1.

### 2.7 Isothermal titration calorimetry (ITC)

Experiments were performed at 25°C using Microcal PEAQ-ITC (Malvern Panalytical, UK). The sample cell was filled with the protein, and the reference cell was filled with water. To minimize heat changes resulting from solution mismatches during mixing, NAD^+^ and NMN ligands were dissolved in the protein buffer solution and drawn into a stirring (titration) syringe for titration. The titration protocol involved injecting 0.4 μL small molecular to 300 μL protein solution for the first point and 2 μL for each of the following 18 points. Dropping one drop takes 4 s, with a 150 s interval between every two drops. The power reference value was 10 cal/s, and the stirring speed was 750 rpm. Prior to the actual experiment, a control experiment was performed using ligands titration buffer to account for the heat change caused by buffer mismatch. The control data should be subtracted to obtain the true binding energy between protein and ligands. Experimental data were fitted to a one-site binding model using Origin scientific plotting software equipped with the instrument. Binding constant (KD) and enthalpy (ΔH) were all flexible parameters, whereas the free energy (ΔG) and entropy (ΔS) were calculated according to the following equation.

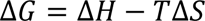

The same ITC experiment was also performed for the DBC1_354-396_ mutants with the E363A, E363K, D372A and D372K mutations.

### 2.8 Molecular Docking

The 3D structures of proteins (wide type and mutants) were obtained from Alphafold2 prediction. The protein receptors were optimized using *AutoDockTools 1.5.6* by adding hydrogen atoms and assigning atomic charges to all protein atoms. Similarly, the two ligands underwent preprocessing, involving charge adjustment, root determination, and torsion angle setting. Afterward, both the protein and ligand are saved as pdbqt format files. *Autodock Vina 1.1.2*[23] was utilized for molecular docking with an exhaustiveness of 12. The binding affinity and interaction between protein and ligands were analyzed using PyMOL[24] and Discovery Studio Visualizer.

## 3. Results

### 3.1 Overexpression and purification of DBC1_354-396_

The construction of DBC1_353-396_ vector is shown in Figure S1A. The 6xHis tag composed of six consecutive histidine residues allows for Ni-NTA affinity chromatography purification[25]. The MBP maintains the solubility of fusion proteins. [26]. The TEV site (ENLYFQ/G) functions as the cleavage site for TEV protease to remove the 6xHis-MBP fusion tag[27]. After cleavage, the target protein retains only two extra glycine (Gly) and alanine (Ala) residues at the N-terminus, minimizing potential effects on its structure and function. The expression and Purification procedure of recombinant DBC1_353-396_ is depicted in Figure S1B. 6xHis-MBP- DBC1_354-396_ fusion protein (49 kDa) and its dimer (98 kDa) were the major products observed after washing with 50mM imidazole (Figure 1A). Nearly 100% of the fusion protein was cleaved after incubation with TEV protease for 3 h, and further purification through Ni-NTA affinity purification yielded a clean and single protein band (Figure 1B). Gel filtration was employed for additional purification and buffer exchange, resulting in a single peak of DBC1_354-396_ (Figure 1C). SDS-PAGE analysis confirmed that this peak exhibited a single band, indicating the high purity of the sample (Figure 1C). In summary, through the aforementioned protein expression and purification methods, we had successfully obtained a highly pure protein. The next step will involve validating the activity of this protein.

**Figure 1.**
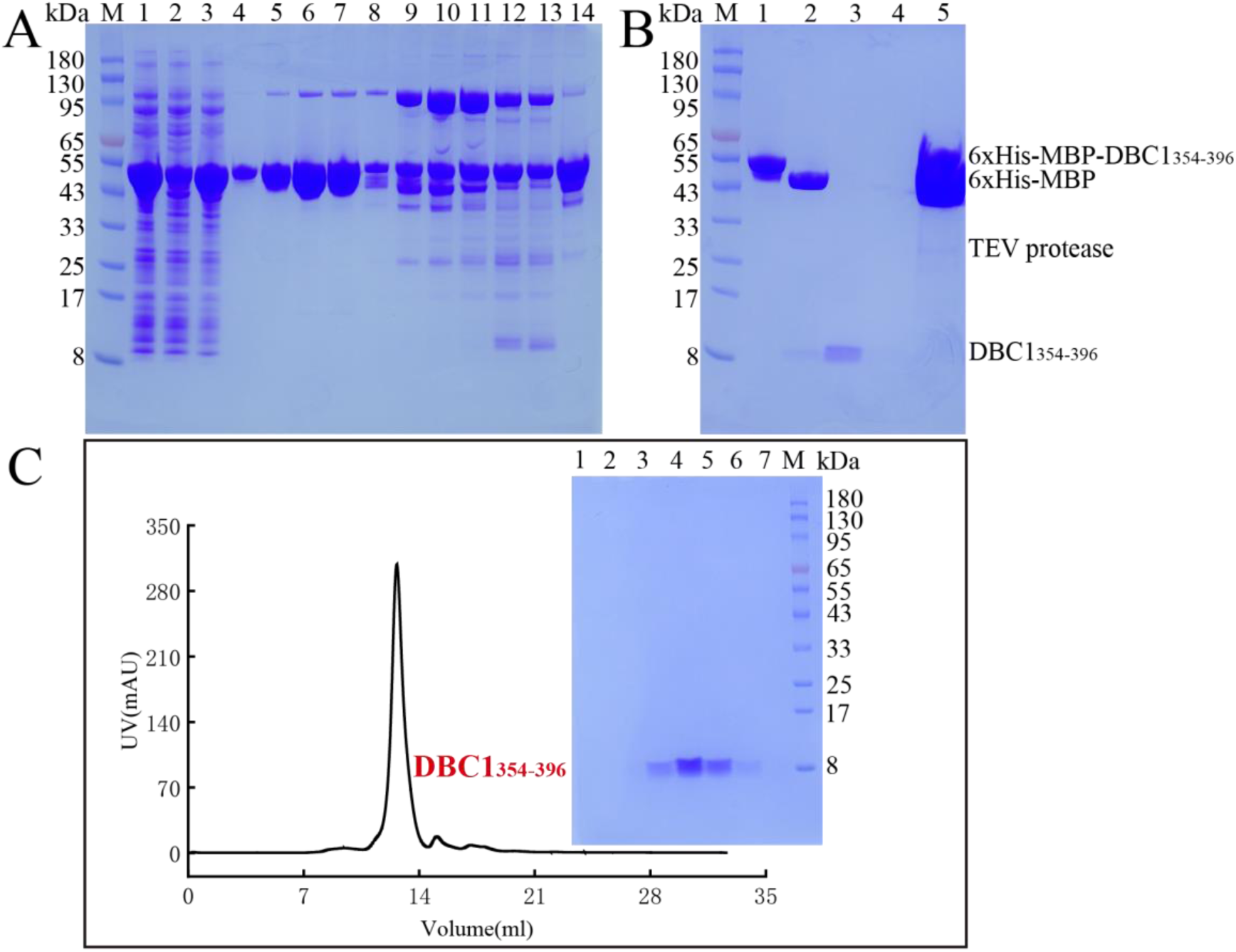
Expression and purification of DBC1_354-396_. (**A)** SDS–PAGE of the samples after Ni-NTA column chromatography. lane 1, pre-column; lane 2-3, flow-through of the lysate after injection through Ni-NTA column; lane 4, 20mM imidazole; lane 5-7, 50 mM imidazole; lane 8-13, 50-1000 mM imidazole; lane 14, lane 10’reduced form. (**B)** SDS–PAGE of the TEV protease cleavage. lane 1, fusion protein without addition of TEV protease; lane 2, 3 h after TEV protease cleavage; lane 3, the flow-through containing DBC1_354-396_ after Ni-NTA column chromatography; (**C)** Spectrogram and SDS–PAGE of pure DBC1_354-396_ after gel filtration column.

### 3.2 HSQC of DBC_1354-396_ and its backbone assignments

HSQC (Heteronuclear Single Quantum Coherence) correlates covalently bonded 1H nuclei with 13C or 15N nuclei, generating cross-peaks. The ^15^N-^1^H HSQC spectrum is the best feedback for identifying whether a protein sample is correctly folded and whether it interacts with other molecules, it can provide information on intermolecular interactions accurate down to the level of amino acid residues or even individual atoms [28-30]. Well-folded and active proteins typically exhibit HSQC spectra with broad width, regular peak shapes, well-dispersed and stable. Here, we obtained the ^15^N-^1^H HSQC spectrum of DBC1_354-396_ at pH 6.5, and the spectrum displayed excellent resolution and separation, indicating that the sample has the correct folding of its 3D structure (Figure 2A). A series of triple resonance experiments successfully assigned the peaks corresponding to each amino acid residue (Figure 2A). The first one or two residues of a protein or peptide are usually not detected due to fast solvent exchange[31]. In a traditional ^15^N-^1^H HSQC spectrum, proline residues often do not exhibit distinct peaks due to lacking backbone amide protons[32]. Accordingly, 42 amide backbone cross peaks with proton frequencies between 7.7 and 8.7 ppm are observed for DBC1_354-396_ (Figure 2A). These data indicated all detectable amino acid residues were successfully scanned by NMR, and the protein sample was well-folded, active, and suitable for titration experiments.

**Figure 2.**
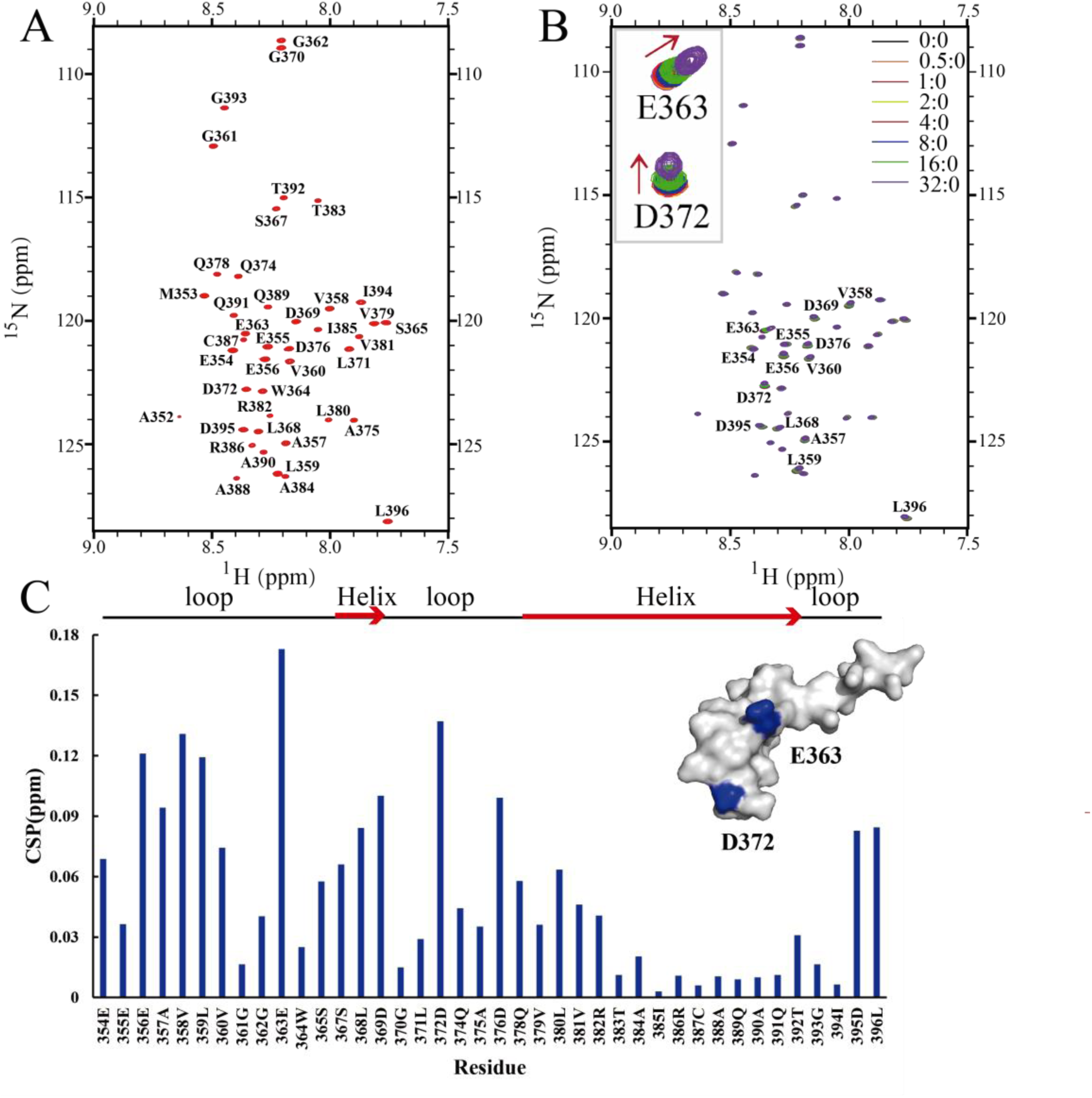
^15^N–^1^H HSQC of DBC1_354-396_. (**A)** ^15^N–^1^H HSQC spectra of DBC1_354-396_ and its backbone Assignment. **(B)** Overlap of ^15^N-^1^H HSQC spectra of titrated-DBC1_354-396_ protein with different concentrations of NAD^+^. **(C)** chemical shift perturbation (CSP) of each amino acid residue when NAD^+^ bind to DBC1_354-396_.

### 3.3 NMR titration revealed the key binding sites

NMR titration has unique advantages in studying the binding mechanism between proteins and ligands, as it can provide information about the binding sites of the ligands with the protein[33]. Ligand binding will change the chemical environment near the binding site, which will induce the chemical shift perturbations (CSP) of affected residues. Such changes can be observed in the ^15^N-^1^H HSQC spectrum[34]. Therefore, we utilized NMR titration to identify the key sites of DBC1_354-396_ binding to NAD and NMN. During the titration of NAD^+^ to DBC1_354-396_, significant chemical shift changes were observed for certain amino acids as the concentration ratio of NAD^+^ increased from 0:1 to 32:1, and a similar effect was seen for NMN (Figure 2B, Figure S2A). This suggests that NAD^+^ and NMN have interactions with DBC1_354-396_. The CSP analysis (Figure 2C, Figure S2B) showed that many amino acids undergo significant chemical shift changes, with E363 and D372 experiencing the most pronounced alterations. This suggests that these two amino acids have the most prominent interactions with the ligands, particularly E363, which exhibits the strongest interaction. Furthermore, NAD^+^ induces a significantly greater CSP compared to NMN, indicating that NAD^+^ has a slightly stronger binding affinity to the protein than NMN. This is consistent with the research findings of Jun Li[7]. These results suggested that NAD^+^ and NMN can bind to DBC1_354-396_, with a higher binding affinity observed for NAD^+^, and the critical binding sites are identified as E363 and D372.

### 3.4 ITC determined the binding affinity of ligands to DBC1_354-396_

After identifying the key sites for ligand-protein binding through the previous experiments, we performed an ITC to go over the binding affinities involved. ITC is a highly sensitive technique used to investigate protein-ligand interactions and assess global changes[35]. Heat is liberated or absorbed as a result of the redistribution of non-covalent bonds when the interacting molecules go from the free to the bound state[36]. ITC measures this heat change and generates a titration curve (kcal/mol vs. molar ratio of ligand/sample), enabling us to determine the binding constant (KD)[37]. When KD >10^-8^ M, it indicates a high-affinity binding between the protein and ligand. When 10^-8^ M < KD < 10^-4^ M, the affinity is considered moderate, while for KD values below 10^-4^ M, the affinity is low. The obtained ITC isotherm for the DBC1_354-396_-NAD^+^/NMN system, clearly demonstrated the spontaneous nature of the binding (Figure 3, Figure S3). Consistent with previous NMR results, NAD^+^ exhibited a moderate binding to DBC1_354-396_, with a measured KD of 8.99e^-6^ ± 8.20e^-6^ M, whereas the binding to NMN was about 2-fold weaker with a KD of 17.0e^-6^ ± 2.30e^-6^ M (Table 2). These results support the hypothesis that NAD^+^ binds to DBC1_354-396_ more tightly than does NMN. Ross’s thermodynamic principles suggest that ΔH ≥ 0 and ΔS > 0 indicate hydrophobic interactions, ΔH < 0 and ΔS > 0 indicate electrostatic interactions, and ΔH < 0 and ΔS < 0 indicate van der Waals forces or hydrogen bonding[38]. Therefore, we can speculate that the forces involved in the binding of NAD^+^, NMN to DBC1_354-396_ are van der Waals forces or hydrogen bonding.

**Figure 3.**
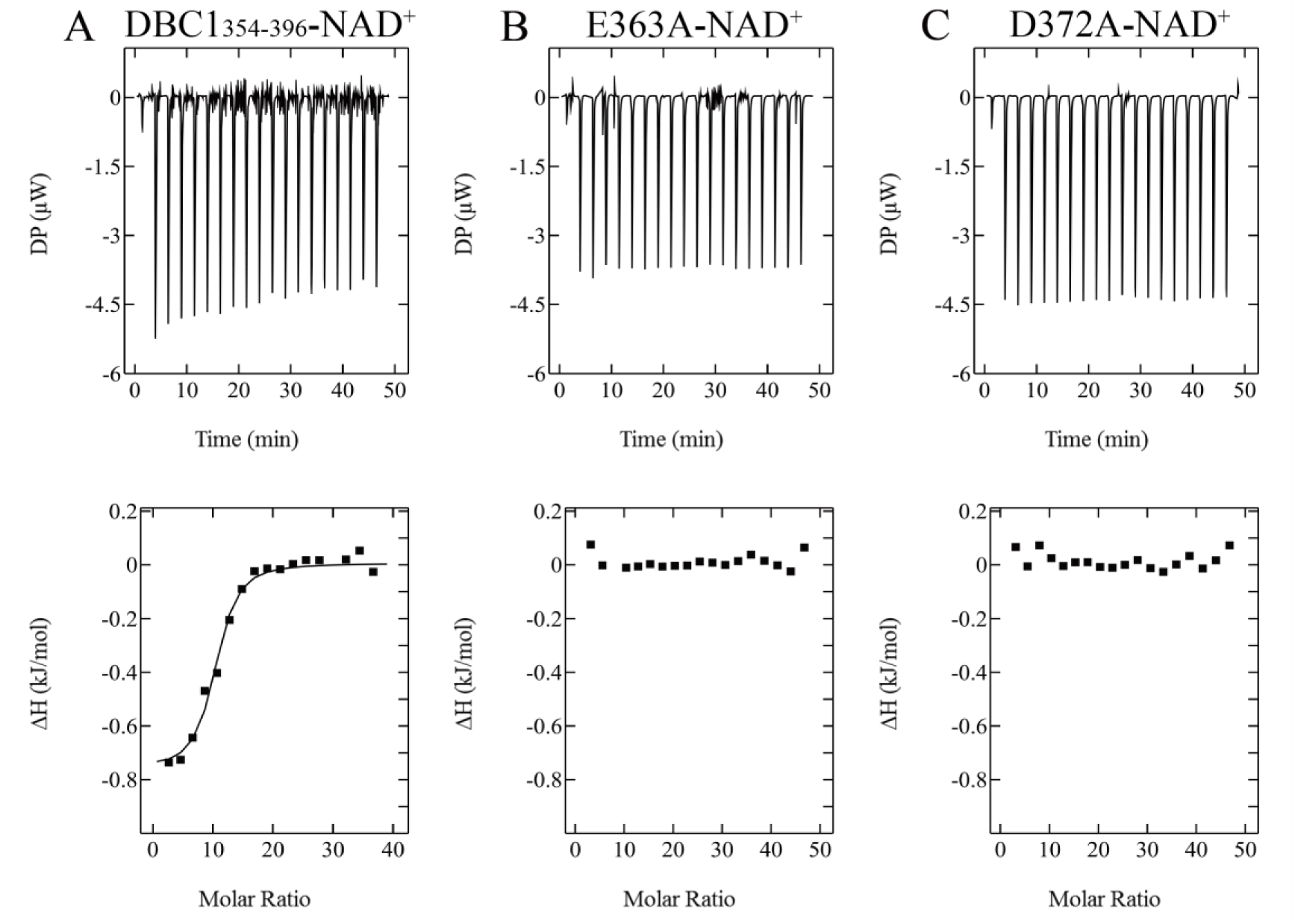
Isothermal titration calorimetry (ITC). (**A)** ITC of DBC1_354-396_- NAD^+^. (**B, C)** ITC of mutant proteins - NAD^+^.

**Table 2.** Thermodynamic parameters obtained for DBC1_354-396_ to NAD^+^/NMN from ITC study.

| Groups | Acquisition errors | Step-down latency (s) | Retention errors |
| --- | --- | --- | --- |
| DBC1<sub>354-396</sub>: NAD<sup>+</sup> | $8.99\text{e}^{-6} \pm 8.20\text{e}^{-6}$ | -28.8 | $-0.767 \pm 9.9\text{e}^{-2}$ |
| DBC1<sub>354-396</sub>: NMN | $17.0\text{e}^{-6} \pm 2.30\text{e}^{-6}$ | -27.3 | $-0.193 \pm 3.7\text{e}^{-3}$ |

### 3.5 ITC of ligands to DBC1 mutants Identified key binding sites

The NMR titration results reveal the critical involvement of E363 and D372 amino acids in the protein-small molecule binding. To gain further insights, we conducted genetic mutations on these key amino acids, employing a principle of altering charged amino acids into uncharged ones or introducing mutations with opposite charges. Such modifications have the potential to influence the protein’s charge distribution, consequently impacting its structure and function, particularly concerning protein-ligand interactions[39]. Therefore, we performed two sets of mutations (E363A, E363K, and D372A, D372K) and employed ITC to assess the binding affinity between the mutated proteins and the small molecules (Figure 3, Figure S3). Remarkably, the results indicated that the heat released upon binding of the mutated proteins to both ligands closely resemble that observed in the control experiment. This suggested an almost negligible interaction between the mutated proteins and the ligands, thereby underscoring the indispensable role of these two amino acids in the binding process.

### 3.6 Molecular Docking predicted binding conformation

Molecular docking is a computational method to predict how a ligand binds within a receptor’s binding site using various algorithms and scoring functions. It shows promise in discovering new drug candidates and understanding protein-ligand interactions[40]. In our study, we identified important binding sites (E363 and D372) and used molecular docking to explore how these protein residues interact with NAD^+^ and NMN. The docking analysis showed that both NAD^+^ and NMN have moderate binding affinities to DBC1_354-396_, with estimated binding energy of -4.9 kcal/mol and -4.5 kcal/mol, respectively (Figure 4, Figure S4). This suggested that NAD^+^ has a stronger affinity to DBC1_354-396_ than NMN, supporting the idea that NAD^+^ binds more tightly to DBC1_354-396_ compared to NMN. The binding patterns of NAD^+^ and NMN with DBC1_354-396_ were shown in Figure 4 and Figure S4. These results indicated that conventional hydrogen bonds and carbon-hydrogen bond play significant roles in the binding process. Specifically, NAD^+^ predominantly forms hydrogen bonds with E363 (2.38 Å) and D372 (2.66 Å), which aligns with the findings from isothermal titration calorimetry (ITC) results (Figure 3). Similarly, NMN interacts with E363 (3.44Å) through a carbon-hydrogen bond and with D372 (2.14 Å) via a hydrogen bond, which also agrees with the ITC data (Figure S3).

**Figure 4.**
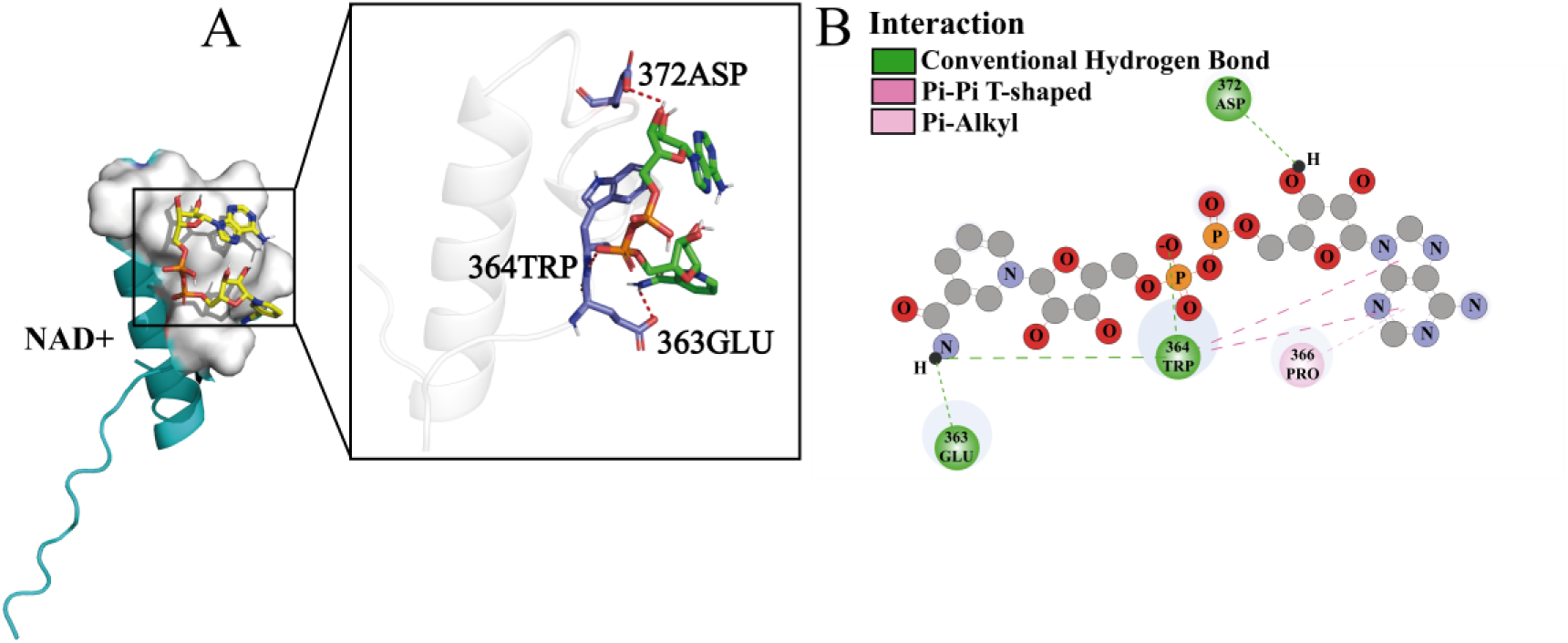
Molecular docking about NAD+ - DBC1_354-396_. **(A)** conformation view of NAD^+^ bind to DBC1_354-396_. **(B)** 2D structural representation of DBC1_354-396_ residues interacting with NAD^+^.

## 4. Discussion

DBC1 (deleted in breast cancer 1) is a large multidomain human nuclear protein that modulates the activities of various proteins. The largest central conserved globular domain in DBC1 is an inactive version of the Nudix hydrolase (MutT) fold, and is likely to bind NAD^+^ derivatives such as ADP ribose[41]. Notably, a study revealed that NAD^+^ binds to the NHD domain of DBC1 without necessitating NAD^+^ cleavage or covalently attached ADP-ribose. This interaction disrupts DBC1-PARP1 complexes in aged mice, rejuvenating PARP activity and reducing DNA damage[7].

To better understand the mechanism by which NAD^+^ binds to DBC1, we initially optimized the DBC1_354-396_ protein’s expression and purification methods, resulting in a highly pure and active protein (Figure 1, Figure S1). Subsequently, a combination of NMR, ITC, and genetic mutation was employed to verify critical binding sites and binding affinity. The results from NMR titration (Figure 2, Figure S2) demonstrated that both NAD^+^ and its precursor NMN interact with the protein, and E363 and D372 are crucial binding sites. Furthermore, the binding affinity of NAD^+^ is superior to that of NMN. The ITC results (Figure 3, Figure S3) confirmed NMR’s findings, showing interactions between both ligands and the protein, with NAD^+^ binding approximately twice as strongly as NMN. Enthalpy-entropy analysis suggested hydrogen bonds drive these interactions. Mutational ITC results revealed mutations hinder small molecule binding, highlighting E363 and D372’s essential role as binding sites. For a better grasp of the roles played by the two crucial binding sites in the binding process, we utilized molecular docking to predict binding conformations and free energies. The findings suggest NAD^+^ mainly interacts with the two key binding sites through hydrogen bonding, while NMN primarily forms hydrogen bond and carbon-hydrogen bond (Figure 4, Figure S4).

The ability of a ligand to bind and modify the activity of a protein target at a specific site greatly impacts the success of drugs in the pharmaceutical industry. One of the most important tools for evaluating these interactions has been high-field solution NMR because of its unique ability to examine even weak protein-drug interactions at high resolution[28, 34, 42, 43]. Isothermal titration calorimetry (ITC) is a complementary technique that can quantify the thermodynamic parameters of intermolecular interactions in situ[44, 45]. Based on these two key methods, this study provides a comprehensive understanding of the affinity and molecular binding mechanisms between NAD^+^/NMN and DBC1_354-396_. It not only lends theoretical backing to NAD^+^-regulated DNA repair but also furnishes valuable direction for advancing targeted therapies and drug research aimed at DBC1.

## 5. Conclusion

This work extensively investigated the molecular mechanism of the binding interaction between DBC1_354-396_ within the NHD region of DBC1 and NAD^+^, revealing the essential binding sites and mode of interaction. This not only establishes a theoretical foundation for unraveling the mechanisms underlying NAD^+^ regulation of protein-protein interactions, but also provides guiding principles for the development of targeted therapies and drug research directed at DBC1. However, it’s important to note that the results of this study are confined to in vitro experiments. Further research and exploration are required to determine whether NAD^+^ and its precursors bind to DBC1 in vivo or even within the human body, contributing to DNA repair and cancer prevention.

