## Supplementary Materials for "Molecular mechanism of NAD^+^ binding to the Nudix homology domains of DBC1"

### Index

**Figure S1** **(A)** Construction of $DBC1_{353\text{-}396}$ vector. **(B)** Flowchart for $DBC1_{354\text{-}396}$ expression and purification.

**Figure S2** **(A)** Overlap of $^{15}N$-$^{1}H$ HSQC spectra of titrated-$DBC1_{354\text{-}396}$ protein with different concentrations of NMN. **(B)** chemical shift perturbation (CSP) of each amino acid residue when NMN binds to $DBC1_{354\text{-}396}$.

**Figure S3** Isothermal titration calorimetry profile of $DBC1_{354\text{-}396}$- NMN, and mutant proteins - NMN/$NAD^{+}$.

**Figure S4** Molecular docking about NMN - $DBC1_{354\text{-}396}$. **(A)** conformation view of NMN binds to $DBC1_{354\text{-}396}$. **(B)** 2D structural representation of $DBC1_{354\text{-}396}$ residues interacting with NMN.

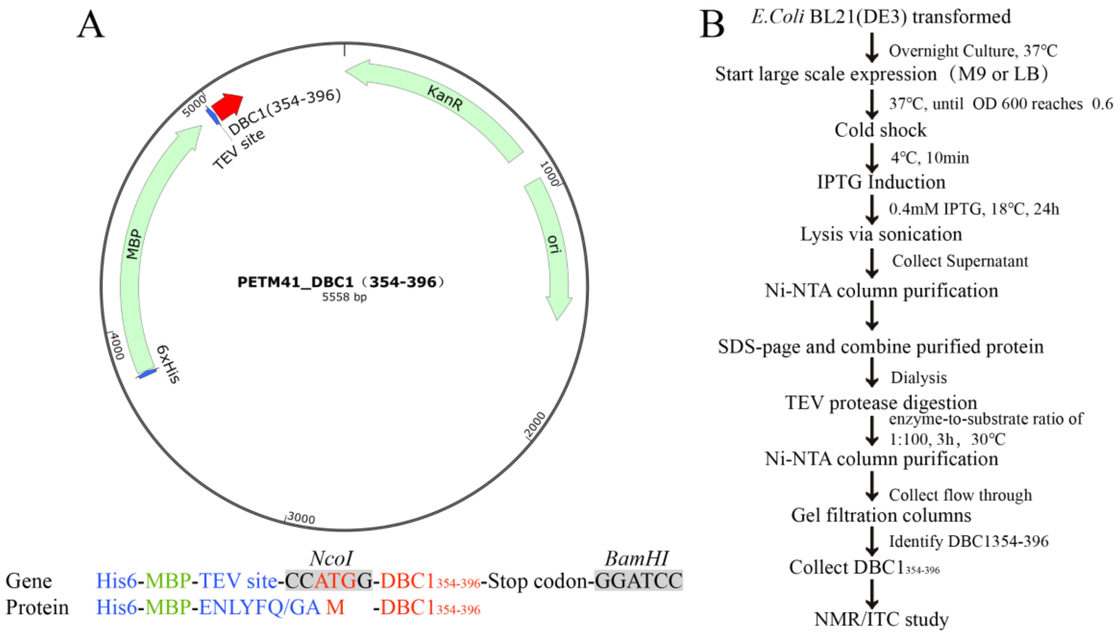
A
KanR
ori
1000
2000
3000
4000
5000
MBP
6xHis
TEV site
DBC1(354-396)
PETM41_DBC1（354-396）
5558 bp
NcoI
BamHI
Gene His6-MBP-TEV site-CCATGG-DBC1354-396-Stop codon-GGATCC
Protein His6-MBP-ENLYFQ/GA M -DBC1354-396
B
E.Coli BL21(DE3) transformed
Overnight Culture, 37℃
Start large scale expression（M9 or LB）
37℃, until OD 600 reaches 0.6
Cold shock
4℃, 10min
IPTG Induction
0.4mM IPTG, 18℃, 24h
Lysis via sonication
Collect Supernatant
Ni-NTA column purification
SDS-page and combine purified protein
Dialysis
TEV protease digestion
enzyme-to-substrate ratio of
1:100, 3h，30℃
Ni-NTA column purification
Collect flow through
Gel filtration columns
Identify DBC1354-396
Collect DBC1354-396
NMR/ITC study

Figure S1

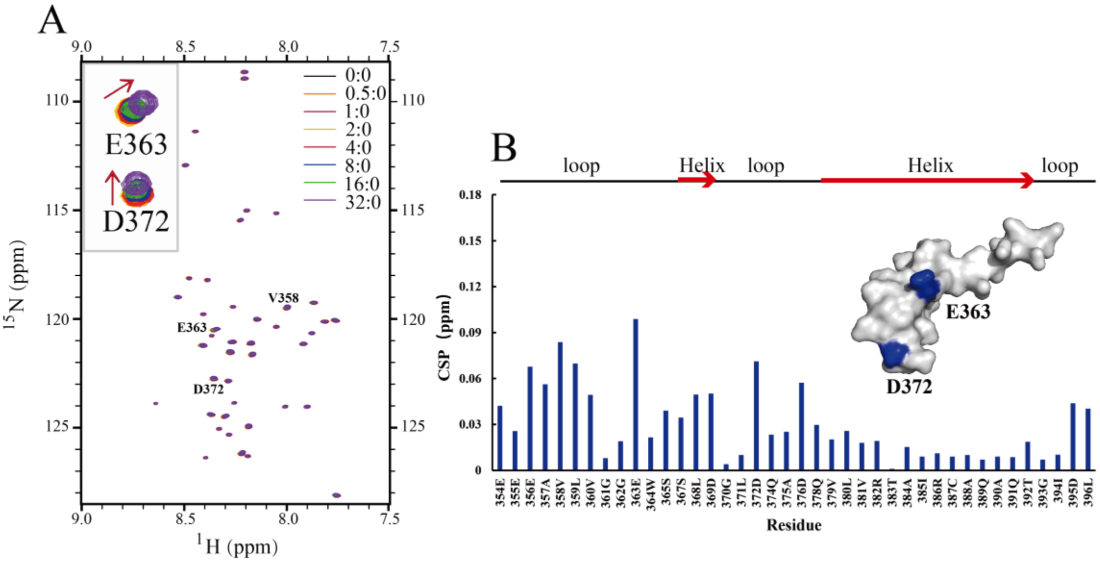
A
B
E363
D372
V358
0:0
0.5:0
1:0
2:0
4:0
8:0
16:0
32:0
$^{15}$N (ppm)
$^{1}$H (ppm)
loop
Helix
loop
Helix
loop
CSP (ppm)
Residue

Figure S2

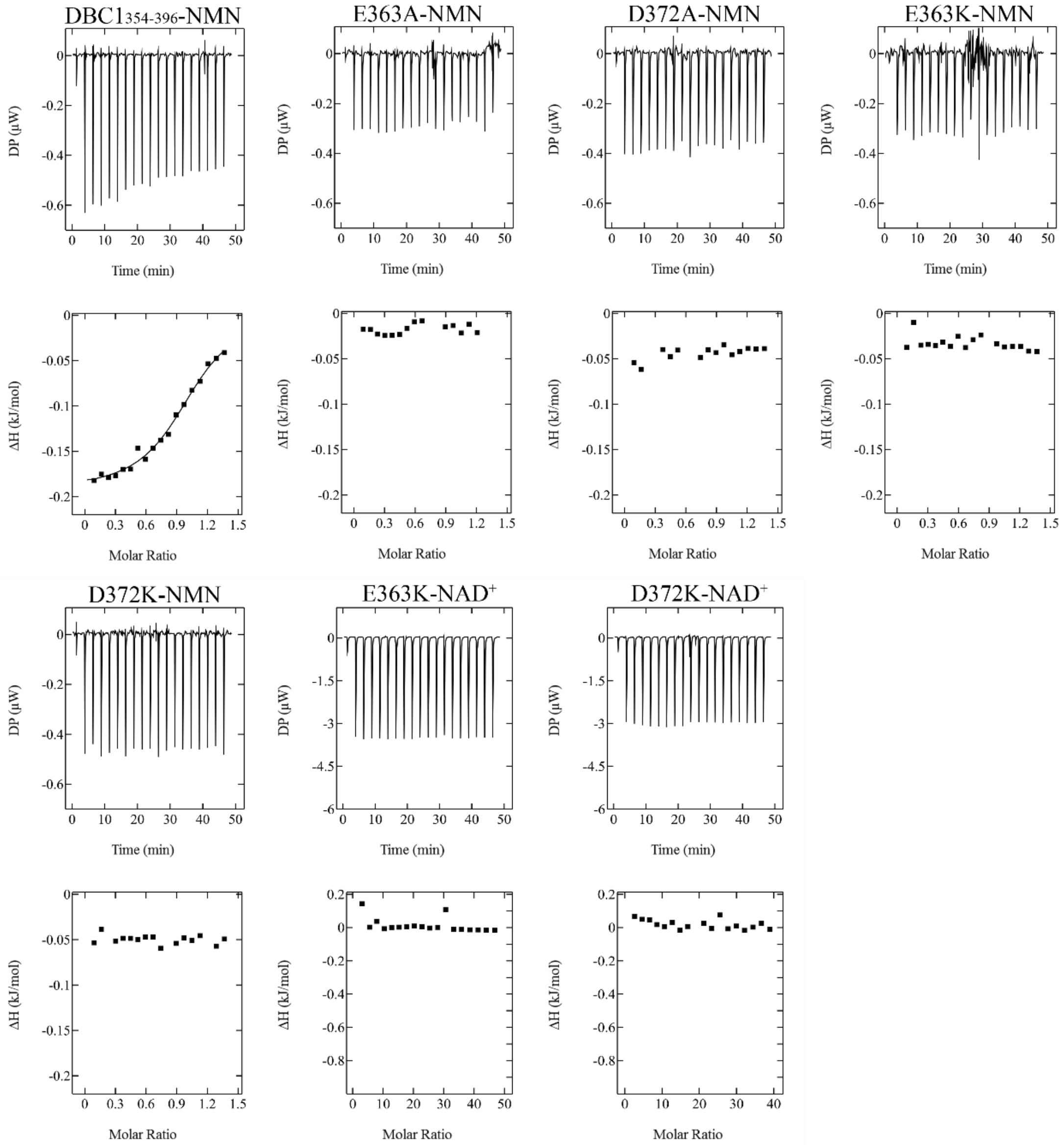

DBC1354-396-NMN
E363A-NMN
D372A-NMN
E363K-NMN
D372K-NMN
E363K-NAD+
D372K-NAD+
DP (μW)
Time (min)
ΔH (kJ/mol)
Molar Ratio


Figure S3

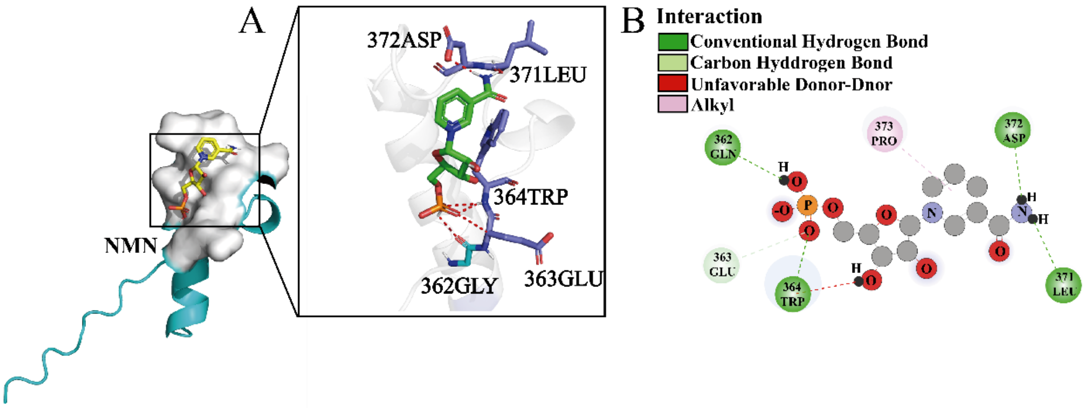
A
372ASP
371LEU
364TRP
362GLY
363GLU
NMN
B
Interaction
Conventional Hydrogen Bond
Carbon Hyddrogen Bond
Unfavorable Donor-Dnor
Alkyl
362 GLN
373 PRO
372 ASP
363 GLU
364 TRP
371 LEU

Figure S4
